# Scaling of ventral hippocampal activity during anxiety

**DOI:** 10.1101/2023.12.22.573072

**Authors:** Carlo Cerquetella, Camille Gontier, Thomas Forro, Jean-Pascal Pfister, Stéphane Ciocchi

## Abstract

The hippocampus supports a multiplicity of functions, with the dorsal region contributing to spatial representations and memory, and the ventral hippocampus (vH) being primarily involved in emotional processing. While spatial encoding has been extensively investigated, how the vH activity is tuned to emotional states, e.g. to different anxiety levels, is not well understood. We developed an adjustable linear track maze (aLTM) for mice with which we could induce a scaling of behavioral anxiety levels within the same spatial environment. Using *in vivo* single-unit recordings, optogenetic manipulations and the application of a convolutional classifier, we examined the changes and causal effects of vH activity at different anxiety levels. We found that anxiogenic experiences activated the vH and that this activity scaled with increasing anxiety levels. We identified two processes that contributed to this scaling of anxiety-related activity: increased tuning and successive remapping of neurons to the anxiogenic compartment. Moreover, optogenetic inhibition of the vH reduced anxiety across different levels, while anxiety-related activity scaling could be decoded using a convolutional classifier. Collectively, our findings position the vH as a critical limbic region that functions as an ‘anxiometer’ by scaling its activity based on perceived anxiety levels. Our discoveries go beyond the traditional theory of cognitive maps in the hippocampus underlying spatial navigation and memory, by identifying hippocampal mechanisms selectively regulating anxiety.

## Introduction

Anxiety is an evolutionary conserved mental and emotional state, underscoring its crucial role in the survival of organisms facing perilous situations (1, 2). Anxiety is characterized by the anticipation of potentially harmful events leading to an attentional bias towards threatening stimuli (3-5). As anxiety prepares an animal or human to face environmental challenges, the level of threat has to be carefully interpreted to support adaptive behavior. An altered perception of negative emotions (6) might contribute to persistent fear and/or chronic anxiety disorders which pose significant personal and societal burdens (7, 8) affecting more than 300 million people worldwide (9). As treatments are available but generic and partially ineffective (10-12), there is an urgent need to understand the neurobiological basis of anxiety in more details. Previous studies have implicated areas of the limbic system including the amygdala, prefrontal cortex and the hippocampus in the generation of emotions (13). But the precise brain region where variations in anxiety levels, and not just anxiety *per se*, are represented and computed remains elusive. We hypothesized that the hippocampus could play an important role in processing different anxiety levels as it is a key structure involved in learning and compares previously stored information (memories) with the current contextual information to generate cognitive maps (14-16). Along the septotemporal or long axis of the hippocampus, differences in synaptic connectivity and molecular expression have been found, inferring functional subdivisions within the hippocampus (17, 18). Studies investigating neural activity patterns and selective lesions of the vH provide evidence, from rodents to its homologue in humans (i.e. the anterior hippocampus), on the role of the vH in the processing of threatening stimuli (19-22). In rodents, anxiety has been frequently studied in mazes such as the elevated plus maze (EPM) which elicit innate anxiety of height and openness as the animals explore the open compartments of the mazes (23-25). vH anxiety neurons, i.e. pyramidal neurons that are activated in the open arms of the EPM, have been linked to anxiety-related behavior through the routing of anxiety-related information to the prefrontal cortex or the lateral hypothalamus (26-28). If and how the activity level of vH neurons relates to the level of perceived anxiety on the EPM is unclear as the precise quantification of emotional states and incentives of animals on an EPM is difficult to assess due to the free exploration and trial- free nature of the task. Furthermore, the specific spatial compartments are inherently linked to the emotional states. Because of such spatial embedding of non-spatial information in many such tasks, there is an ongoing discussion on how much non-spatial information, such as emotions, are actually defined by a spatial framework or can be encoded independently in the hippocampus (29, 30).

To address the nature of anxiety coding in the vH, we developed a linear track maze (31) that could be gradually adjusted to elicit different anxiety levels to an open and elevated compartment within the same spatial environment. With this fully automated adjustable linear track maze (aLTM), we could overcome some of the rigidities and difficulties of the EPM allowing us to define trials and achieve a scaling of anxiety levels while recording and manipulating the activity of neurons in the vH.

## Materials and Methods

### Subjects

Male C57BL6/J mice (N = 28) aged 4-6 months (Janvier Labs, France) were group housed with *ad libitum* access to food and water in a temperature-controlled room on a 12-hour light/dark cycle. Following surgery, mice recovered for at least 21 days. Behavioral experiments were conducted in the light phase and mice were food deprived to reach 90% and not less than 85% of the baseline weight (= average weight of 3 consecutive days prior to the experiment day). All experimental protocols adhered to the guidelines set forth by the Animal Welfare Office at the University of Bern and received approval from the Veterinary Office of the Canton of Bern.

### Surgical procedures

18 mice were anesthetized with isoflurane (Attane, Provet, induction 4% in oxygen at a flow rate of 1L/min in anesthetic box, maintenance of the anesthesia based on 1.5-2% of isoflurane in oxygen at a flow rate of 0.8 L/min during the entire surgery) and body temperature was maintained at 37°C with a temperature controller (Harvard Apparatus). Analgesia was administered upon induction and if needed for 3 days post-operation (Carprofen, 2-5 mg/kg subcutaneous). The incision site was infiltrated with lidocaine (2%, Sintetica). Implants were secured to the skull using light-cured dental adhesive (Kerr, OptiBond Universal) and dental cement (Ivoclar, Tetric EvoFlow). After surgery, mice were returned to their home cages for recovery and kept warm for 3 days by warming the cages through a heating pad (Medisana, HP625).

### Virus injection and fiber implantation

For optogenetics experiments, 12 mice were infused bilaterally in the vH with the following coordinates: AP: -3.08 mm, ML: ±3 mm, DV: -3.8 mm. The viral solution (250 nL) was delivered via a glass micropipette (30-40 μm tip) attached by tubing to a Picospritzer III microinjection system (Parker Hannifin Corporation) at a rate of 25 nL/min. Bilateral infusion of either ssAAV5-CamKIIα-eArchT3.0-EYFP or ssAAV5-CamKIIα-EYFP (ETH Zurich Viral Vector Facility) was immediately followed by optic fibers implantation (200 μm core, 0.37 numerical aperture, Thorlabs) above the vH at the following coordinates: AP: -3.08 mm, ML: ±3 mm, DV: -3.75 mm (Figure 2A). The optic fiber implants were fixed to the skull with stainless steel screws, resin-based dental cement (A1 Tetric EvoFlow, Ivoclar Vivadent) and light-cured dental adhesive (Kerr, OptiBond Universal). To ensure sufficient expression of the viral construct a minimum of five weeks was allowed for the mice to recover before conducting the behavioral experiments.

**Figure 1.**
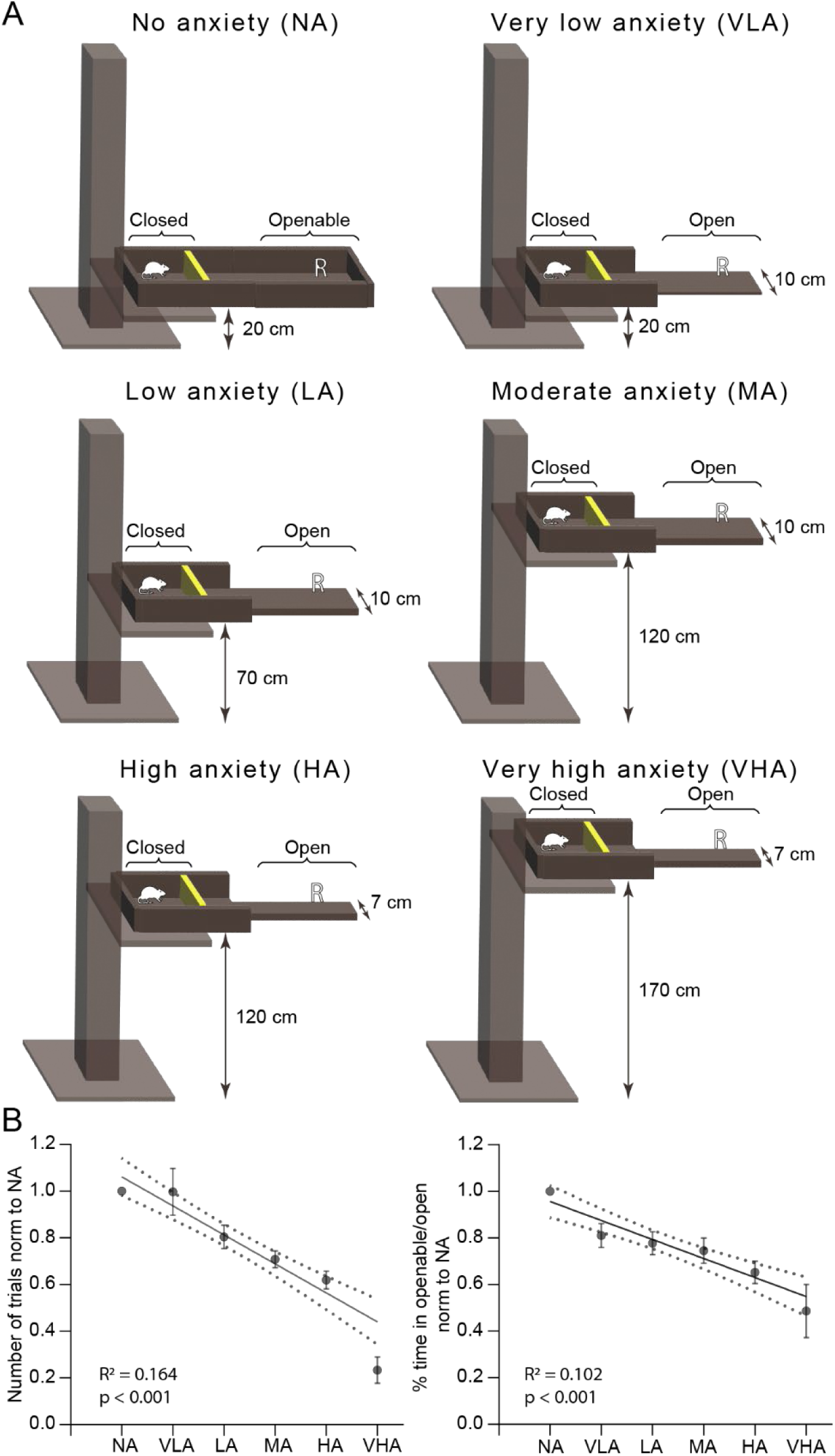
Different configurations of the aLTM induce a scaling in anxiety behavior. (A) aLTM’s anxiety configurations. The yellow rectangle in the maze represents an automated door that closed after each trial for a random amount of time (between 10-40 seconds) to avoid stereotypic behavior. ‘R’ indicates the reward delivery in the openable/open part of the aLTM. **(B)** Number of trials normalized to trials taken in NA (left) and percentage of time in the openable compartment normalized to NA (right) across anxiety configurations indicate gradually increasing behavioral anxiety from NA to VHA. Data represented as linear regressions with 95% confidence intervals calculated based on all behavioral sessions (n = 64 sessions from N = 16 mice,). Means ± SEM of all behavioral sessions for each anxiety configuration are also indicated.

**Figure 2.**
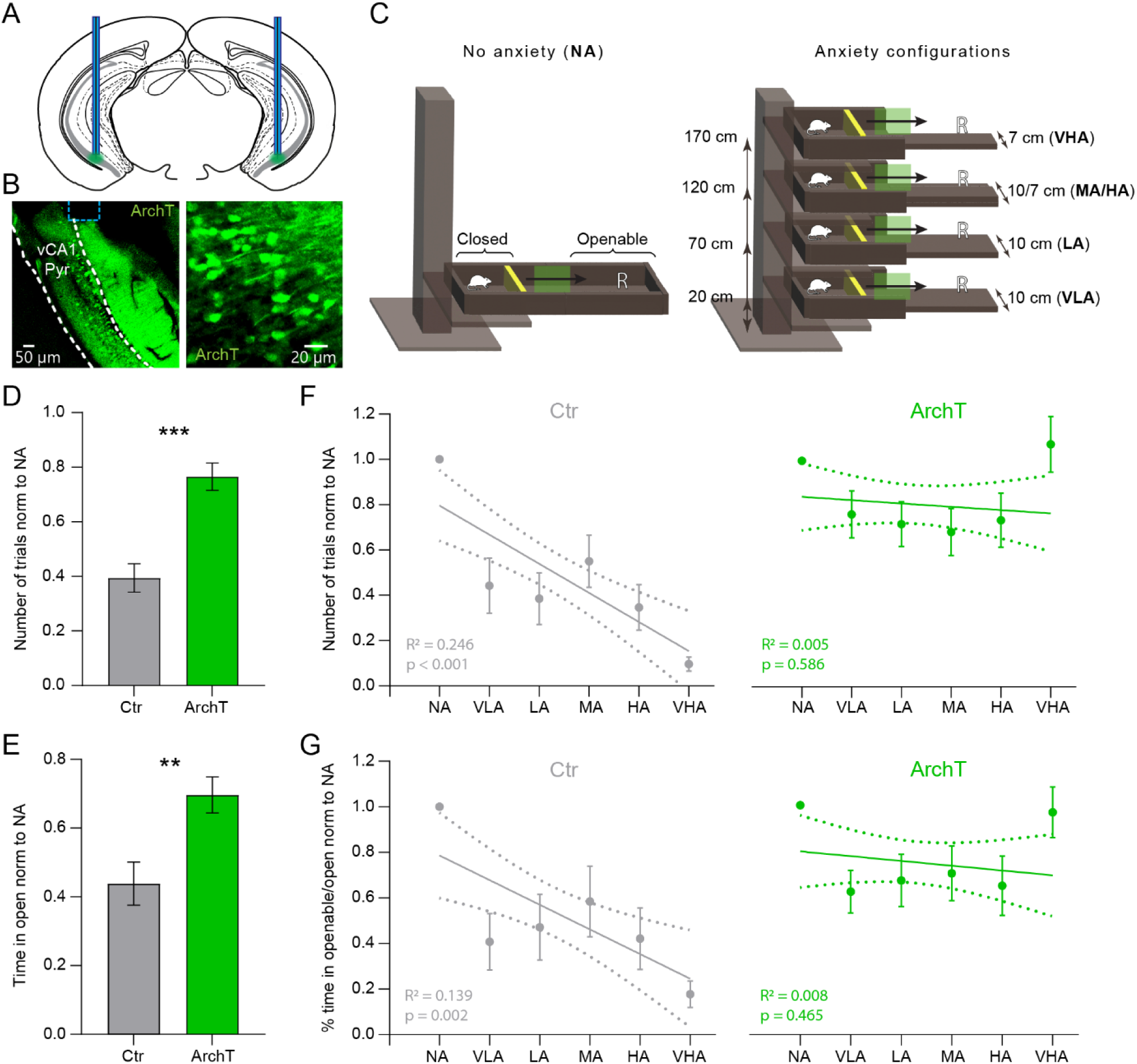
vH inhibition impairs the scaling of anxiety behavior. (A) Schematic representation of optic fiber implants (blue rectangles) and injection sites (green circles) of adeno-associated viruses (AAV5) expressing ArchT and eYFP reporter under the control of a CamKIIα promoter. **(B)** Image of anti-YFP immunolabelling in vH of ArchT mice with a magnification of vH somata on the right. vCA1 Pyr: ventral CA1 stratum pyramidale; Dashed blue line: optic fiber **(C)** Schematic representation of optogenetic intervention in outbound trajectories indicating with a green rectangle the start of light application when the mice reach halfway from the door position to the open space and ending when the mice fully entered the open space. **(D)** ArchT mice (N = 6) ran more trials to the open part of the maze across anxiety configurations (top, Mann-Whitney test *p* <0.001) and **(E)** spent significantly more time in the open part than eYFP control (ctr) mice (N = 6) (bottom, Mann-Whitney test *p* = 0.001). **(F)** Gradually decreasing number of trials taken in the aLTM’s open part were disrupted in ArchT mice (right) but not control mice (left). **(G)** Gradually decreasing time spent in the aLTM’s open part was disrupted in ArchT mice (right) but not control mice (left). Data represented as linear regressions with 95% confidence intervals calculated based on all behavioral sessions (n = 66 sessions). Means ± SEM of all behavioral sessions for each anxiety configuration are also indicated.

### Optogenetic manipulation

For the optogenetic inhibition of CamKIIα^+^ vH neurons during linear track maze navigation, a continuous light pulse was delivered when the mouse transitioned from the closed to the open part of the maze (Figure 2C) via a patch cord coupled to a laser beam (Cobolt 06 – DPL 561 nm, Hübner Photonics). The laser was triggered automatically when the mouse’s body center was detected inside the second half of the transition and was turned off as the mouse’s body center was detected in the openable compartment using the ANY-maze animal tracking software.

### Surgery for *in vivo* electrophysiological recordings

6 mice were implanted with a custom-made microdrive (Axona Ltd.) containing 8 independently moveable tetrodes made of four gold-plated (NanoZ, Multi Channel Systems) twisted tungsten microwires (12.7 μm inner diameter, California Fine Wire Company, impedances < 150 kΩ). Microdrives were lowered to the vH. Implants were fixed to the skull with 5 miniature stainless steel screws (00-96x1/16, Bilaney) with two of them placed above the cerebellum and connected to the electrode interface board to serve as ground for the electrophysiological recordings.

### Adjustable linear track maze (aLTM)

A home-made adjustable linear track maze (31) (L80 cm x W10-7 cm x H12 cm, Suppl. video 1) was built in order to create 6 different anxiety levels in mice: 1. a ‘no anxiety’ (NA, Figure 1A top left) configuration in which the entire maze was surrounded by walls and the mice had no visual insight into the height of the maze. 2. a ‘very low anxiety’ (VLA, Figure 1A top right) configuration in which the openable second half of the maze was opened and the mice could perceive the 20 cm height of the maze. 3. a ‘low anxiety’ (LA, Figure 1A middle left) configuration in which the openable second half of the maze was opened and the mice could perceive the 70 cm height of the maze. 4., a ‘moderate anxiety’ (MA, Figure 1A middle right) configuration where the maze was further elevated by a minilift (ESM100, Dahlgaard & Co.) to 120 cm from the ground. 5. the ‘high anxiety’ (HA, Figure 1A bottom left) configuration in which the maze was still at 120 cm height but the open part was changed to a narrower width from 10 cm to 7 cm. 6. the ‘very high anxiety’ (VHA, Figure 1A bottom right) configuration with the maze further elevated to 170 cm with 7cm width of the open part. 5 mg reward pellets were randomly placed in different distal locations on the openable part after each trial to motivate the mice. A custom made automatic door (Stepper driver ROB-12859, SparkFun Electronics; Cypress Semiconductor Corp; Switch snap, C&K) in the closed part was built in order to break stereotypic behaviors and create a safe environment for resting in-between sessions without any interference from the experimenter (Figure 1A).

Each different anxiety configuration was presented twice with more anxiogenic or less anxiogenic configuration orders at 50 Lux lighting conditions. Scrambled configuration experiments were performed in the following order: LA-VLA-HA-NA-VHA-MA (Suppl. Figure 2). For each experiment the mouse was first placed into the closed compartment with the door closed for 300 seconds for habituation and rest. Then the door was opened and the mouse was free to initiate a trial or not. A trial is considered for every trajectory that the mouse performed going from the first half of the maze to the second half and came back to the first half. If the mouse explored the second half of the maze (the openable compartment) and returned to the closed part, the door would be closed for a random time of 10-40 seconds to break stereotypic locomotion and allow the experimenter to place the reward in the random spot in the second anxiogenic part of the maze. For each configuration, the duration that the mouse could run trials lasted 500 seconds. In a 2-minute break between each configuration, the mice rested in the closed compartment with the door closed. In the novel configuration, a control NA session would be compared to a ‘novel’ configuration (300s rest + 500s with trials) where the walls in the openable part would be replaced by walls decorated with stripes of a different color (yellow stripes) and different texture (tape) (Figure 5A right).

**Figure 3.**
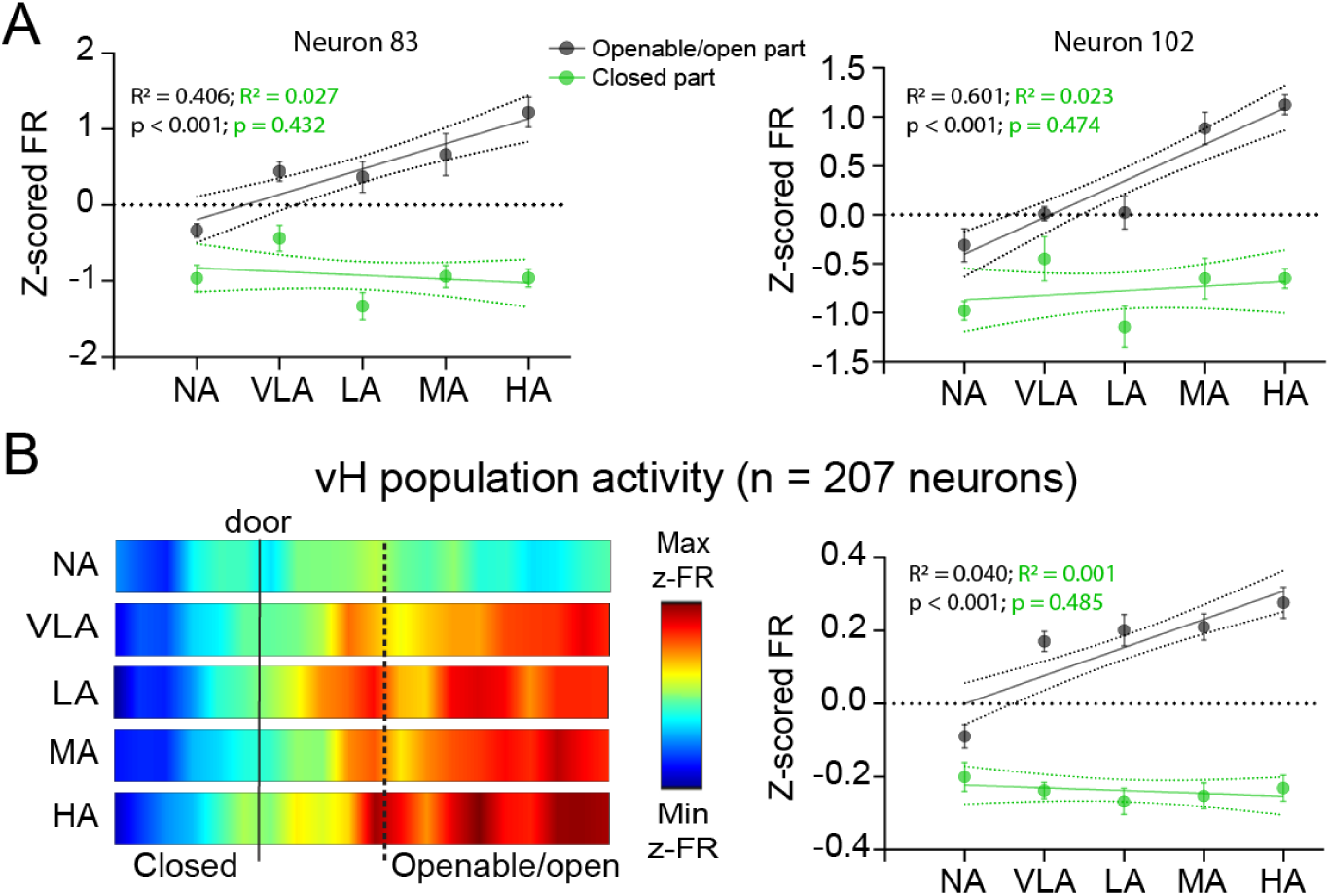
Scaling of vH activity during anxiety. (A) Individual example neurons with increasing z-scored activity in the aLTM’s open part across anxiety levels (linear regression on values of all sessions across NA to HA for Neuron 83 (left) and Neuron 102 (right). **(B)** Left, heatmap with z-scored firing rates show vH population activity progressively increasing in the aLTM’s open part compared to the closed part. Heat maps depict the linearized z-scored firing rate activity of the entire recorded population (N = 6 mice, n = 207 neurons). Right, linear regression on all activity values of all sessions in the open part and in the closed part. Data represented as linear regressions with 95% confidence intervals based on all z-scored firing rate recorded values. Means ± SEM of all behavioral sessions for each anxiety configuration are also indicated.

**Figure 4.**
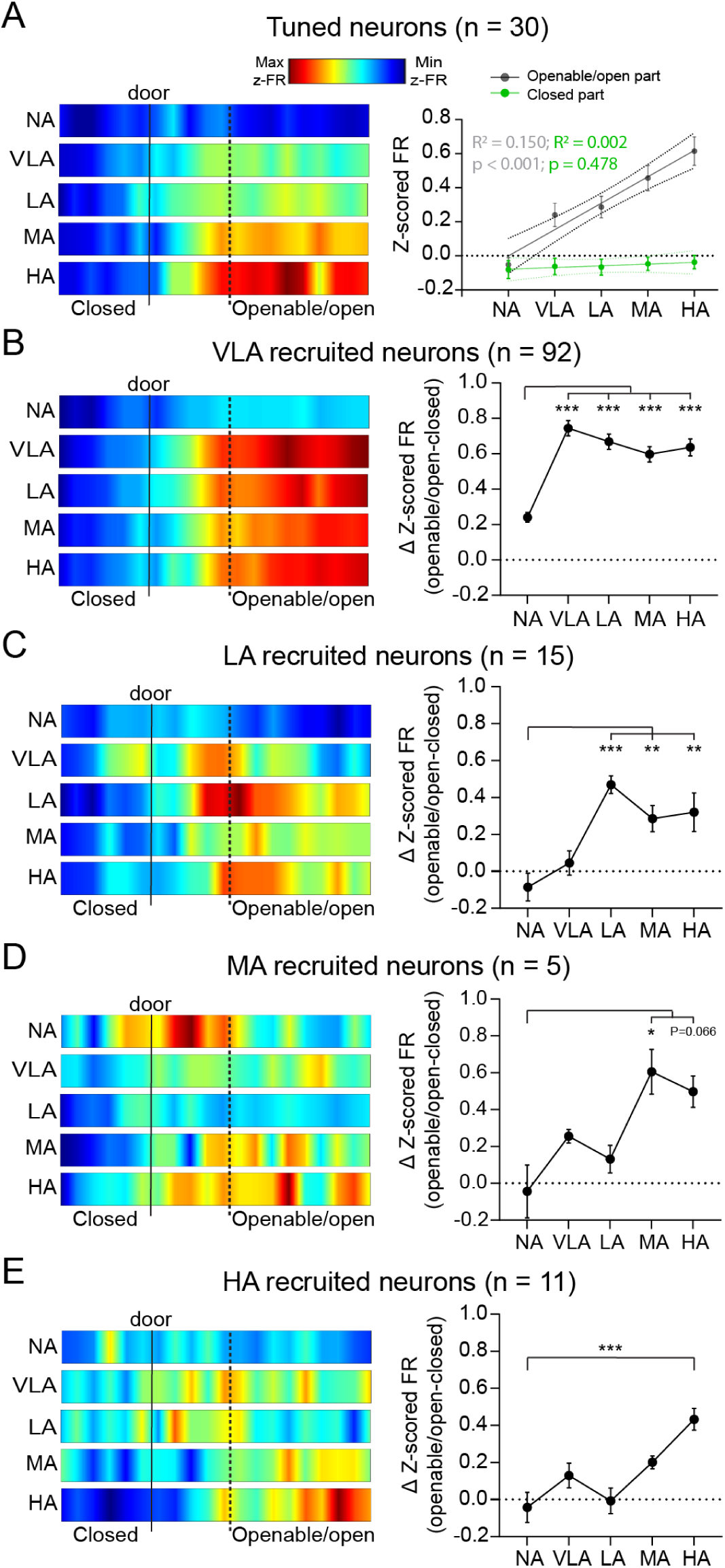
Tuning and reorganization of vH neuronal activity underlies scaling during anxiety. (A) Left, linearized z-scored firing rate heat maps of all neurons showing an increase in activity at each anxiety configuration in the aLTM’s open but not the closed part. Right, linear regressions between z-scored firing rates and anxiety configurations. Data represented as linear regressions with 95% confidence intervals based on all recorded z-scored firing rate values (N = 6 mice, n = 30 neurons). Means ± SEM for each anxiety configuration are also indicated. **(B)** Left, linearized z-scored firing rate heat maps of all VLA recruited neurons. Right, z-scored firing rate differences between open to closed activities across levels in VLA recruited neurons show reorganized neuronal activity at VLA configuration that persists at higher anxiogenic configurations. VLA recruited neurons: Friedman test, *p* < 0.001; Dunn’s multiple comparisons test, NA vs VLA *p* < 0.001; NA vs LA *p* < 0.001; NA vs MA *p* < 0.001; NA vs HA *p* < 0.001 (n = 92 neurons). **(C)** Left, linearized z-scored firing rate heat maps of all LA recruited neurons. Right, z-scored firing rate differences between open to closed activities across levels in LA recruited neurons show reorganized neuronal activity at LA configuration that persists at higher anxiogenic configurations. LA recruited neurons: RM one-way ANOVA, *F*(2.93,40.98) = 13.98, *p* < 0.001; Dunnett’s multiple comparisons test, NA vs VLA *p* = 0.360; NA vs LA *p* < 0.001; NA vs MA *p* = 0.002; NA vs HA *p* = 0.003 (n = 15 neurons). **(D)** Left, linearized z-scored firing rate heat maps of all MA recruited neurons. Right, z-scored firing rate differences between open to closed activities across levels in MA recruited neurons show reorganized neuronal activity at MA configuration that persists at the higher anxiogenic configuration. MA recruited neurons: Friedman test, *p* = 0.007; Dunn’s multiple comparisons test, NA vs VLA *p* > 0.999; NA vs LA *p* > 0.999; NA vs MA *p* = 0.020; NA vs HA *p* = 0.066 (n = 5 neurons). **(E)** Left, linearized z-scored firing rate heat maps of all HA recruited neurons. Right, z-scored firing rate differences between open to closed activities across levels in HA recruited neurons show reorganized neuronal activity at HA configuration. HA recruited neurons: Friedman test, *p* < 0.001; Dunn’s multiple comparisons test, NA vs VLA *p* = 0.319; NA vs LA *p* > 0.999; NA vs MA *p* = 0.236; NA vs HA *p* < 0.001 (n = 11 neurons). N = 6 mice. Dotted line in the heat-maps represents entrance location in the openable part. Error bars represent mean ± SEM.

**Figure 5.**
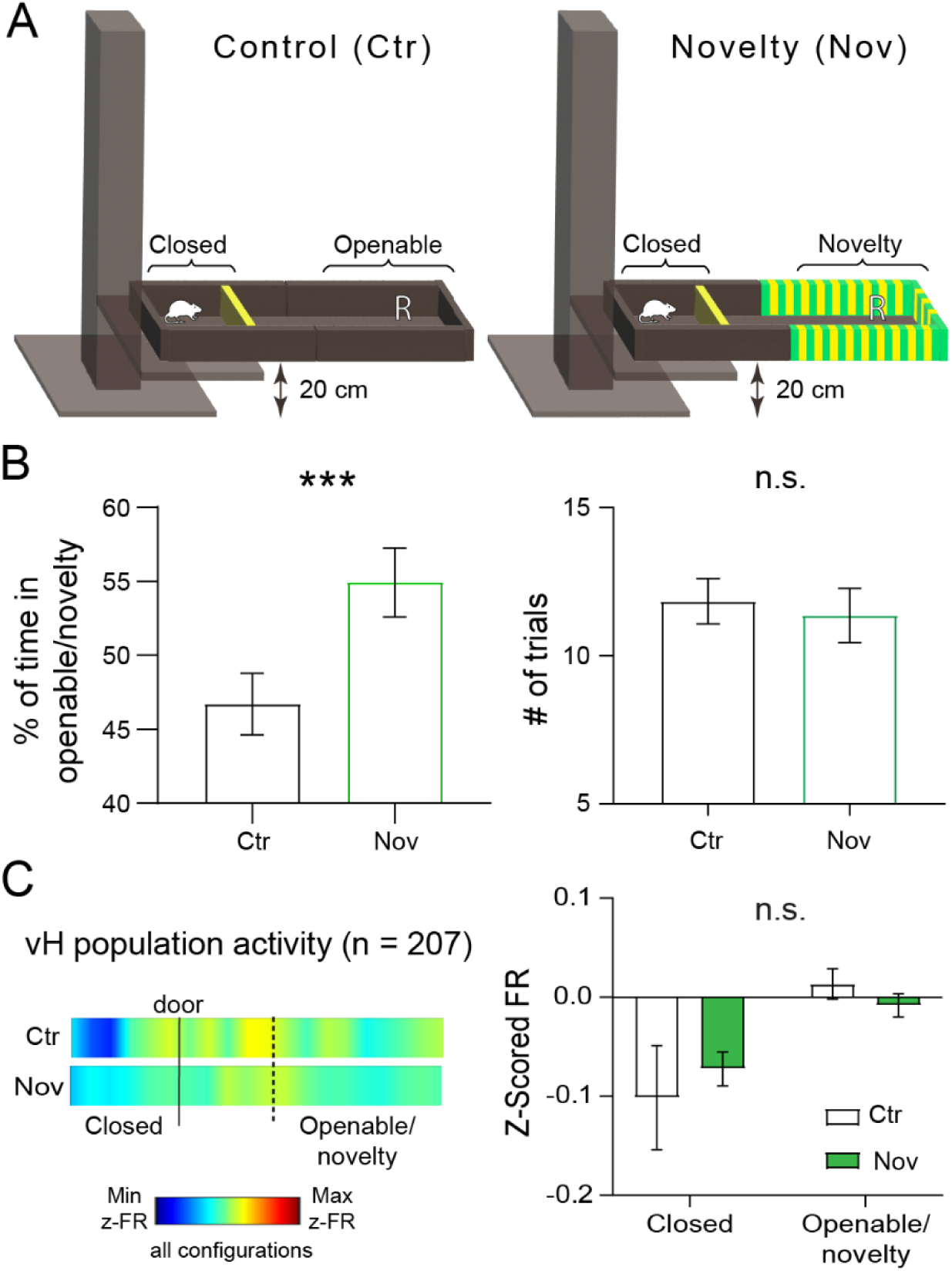
Novelty does not change vH population activity. (A) Control (left) and ‘Novelty’ (right) configurations of the aLTM illustrating the visual differences of the novel part of the maze. **(B)** Mice spent more time (left) in the novel part of the maze but did not engage in different number of trials (right) than in the control configuration. Wilcoxon test, *p* < 0.001. Paired t-test *p* = 0.392. **(C)** Left, linearized firing rate heatmaps and, right, z-scored firing rate show no significant changes induced by the novelty and that activity levels did not reach values close to the maximum vH activations of the anxiety configurations. Two-way RM ANOVA Interaction values: *F*(1,12) = 1.68, *p* = 0.219. Uncorrected Fisher’s LSD: ctr, closed vs open *p* = 0.002; nov, closed vs open *p* = 0.070; closed, ctr vs nov *p* = 0.372; openable, ctr vs nov *p* = 0.373 (N = 6 mice, n = 207 neurons). Dotted line in the heat-map represents the entrance location of the openable part. Error bars represent mean ± SEM.

For anxiety testing, every mouse had been challenged up to 4 times with an interval of at least 5 days between each experiment. Videotaping was carried out from above the maze with the behavior system ANY-maze (www.any-maze.com). Lighting was maintained at 50 lux in each configuration thanks to a dimmable light.

### Histology

Mice were overdosed with a 5% ketamine and 2.5% xylazine solution (0.02 ml/g) and transcardially perfused with phosphate buffer saline (0.1M PBS, pH 7.4, Roche) followed by 4% paraformaldehyde (PFA, Roti Histofix 4 %, Roth). For electrophysiology experiments, mice were anaesthetized with isoflurane (as stated above) prior to perfusion and electrolytic lesions were made at the tip of each recording tetrode by applying 5 times 30 μA for 2 seconds. Brains were kept in 4% PFA for 24 hours post-fixation. After a rinse in PBS for at least 24 hours at 4°C, brains were sectioned coronally at a thickness of 50 μm using a vibratome (VT1000 S, Leica Microsystems) and preserved in 0.5% sodium azide PBS at 4°C. Lesions were located and only tetrodes positioned in the vH were included for further analysis (Suppl. Figure 1).

### Immunohistochemistry

Free floating brain slices were blocked with 5 % normal donkey serum (NDS, Abcam) in 0.1% Triton X-100 in PBS (PBS-T) for 30 minutes on a shaker at room temperature (25 °C). Sections were incubated for 72 hours at 4°C with an anti-GFP antibody (1:100, Abcam, ab13970) washed with PBS-T for 5 minutes (3x) and incubated for 2 hours at room with secondary antibody (AlexaFluor-488 1:100, Jackson ImmunoResearch Laboratories, 703545145). Slices were mounted onto slides (ThermoFisher) and coverslipped with aqueous mounting medium (Aqua-Poly-Mount, Polyscience). Sections were imaged on a stereomicroscope (Leica M205 FCA, Leica) mounted with a monochrome camera (Leica DFC345 FX, Leica) and a UV light source (X-CITE, Lumen Dynamics) or with a confocal microscope (LSM 880, Zeiss, Jena, Germany) and stored at 4°C.

### Behavioral quantification

For behavioral assessments, we measured the number of trials completed by the mice and the percentage of time spent within the openable section of the maze per presented configuration. These measurements were normalized relative to the total number of trials completed and the duration of time spent in the openable maze section during the NA configuration. This normalization accounted for variations in locomotion and exploration tendencies among individual mice.

### Behavior with electrophysiological recordings

After 21 days post-surgery, the 6 mice were familiarized to the experiment room and acclimated to the handling procedures, which included cable tethering to the recording system. On the following day, tetrodes were lowered by roughly 150 μm while monitoring hippocampal electrophysiological hallmarks (i.e. sharp-wave-ripple and theta oscillations amplitude) and multi-unit spiking activity of the stratum pyramidale was detected. Electrophysiological signals were synchronized to behavior via TTL outputs sent from ANY-maze (at 30 Hertz). Neurophysiological data was collected and analyzed from levels NA, VLA, LA, MA, HA but not VHA due to the low sampling of this open part.

### Spike detection and unit classification

The extracellular electrical signals from the tetrodes were amplified, filtered and digitized with a headstage (Intan RHD amplifier, Intan Technologies). The signals were acquired using an Intan RHD2000 evaluation board (Intan Technologies) at a sampling rate of 20 kHz. Neuronal spikes were extracted offline by either using Kilosort2 (63, 64) or by detecting signal amplitudes five standard deviations above the root mean square of the digitally filtered signal (0.8 - 5 kHz) over 0.2 ms sliding windows. A principal component analysis was implemented to extract the first three components of spike waveforms of each tetrode wire (65, 66). Spike waveforms were automatically clustered using KlustaKwik (http://klustakwik.sourceforge.net/). Individual units were manually refined in Klusters (67).

### Tuning and recruitment quantification

A neuron was classified as anxiety-tuned if its firing rate in the open part of the aLTM was significantly correlated (Spearman correlation analysis) with the linearly increasing anxiety configurations. Neuronal recruitment was assessed by comparing the neuron’s firing rate in the open part of the maze to its firing rate in the closed part. The first configuration in which the cell showed a statistically significant increased firing rate in the open part compared to the closed part was used to classify that neuron as being recruited in that specific configuration.

### Computational methods

Calculations were performed on UBELIX (http://www.id.unibe.ch/hpc), the HPC cluster at the University of Bern. For quantification performed on neuronal populations, z-scored neuronal activities were calculated to allow unbiased comparisons of firing rates. Each neuron was independently z-scored based on its activity in the entire recording session.

Decoding of the anxiety level was performed using the ResNet18 architecture (68) distributed as part of PyTorch (69). Networks were trained using a cross-entropy loss with the ADAM optimizer (70) with the following parameters: Learning rate γ=0.01; Weight decay parameter λ=0.0025; Dropout probability p=0.1; Ratio of training data to training and validation data: 70%; Number of epochs: 400; Batch size: 150; Number of time steps: N = 25.

For each training or test sample, the input to the network is an m*N matrix, where m is the number of recorded neurons in the session.

### Statistical analysis

Analyses were conducted in GraphPad PRISM and MATLAB using custom scripts. All datasets were tested for normality using the D’Agostino & Pearson test, and if found to be normally distributed, we used parametric tests or otherwise the equivalent non-parametric tests. In case of multiple comparison, significance was evaluated with a one-way or two-way repeated measures ANOVA or Friedman’s test followed by Dunn’s, Dunnett’s or Uncorrected Fisher’s LSD post hoc tests. To evaluate scaling and linearity induced by the different anxiogenic configurations, we applied Spearman correlation and simple linear regression analyses against a numerical linear increasing vector representing the anxiety levels across which we determined linear scaling of anxiety behavior. All plots show the best-fit line with its R^2^ and *p* values and the 95% confidence bands. Both the Spearman correlation and the simple linear regression calculations were performed on the entire dataset of behavioral or electrophysiological individual values. However, to improve visualization, plots also present the mean ± SEM.

### Data availability

The datasets generated during the current study are available from the corresponding authors upon request. Matlab and Python files are available in the following GitHub repository: https://github.com/CarloCerquetella/aLTM.

## Acknowledgments

This work was supported by the following grants: an ERC starting grant (716761) to S.C., a Swiss National Science Foundation professorship grant (170654) to S.C., a Swiss National Science Foundation grant (31003A_175644) to J.P.P and C.G, and a Swiss National Science Foundation grant (P500PM_210800) to C.G.. C.C. and S.C. designed the experiments. C.C. performed experiments and data analysis. C.C., S.C. and T.F. interpreted the data, prepared and wrote the manuscript. C.G. and J.P.P contributed to data analysis and edited the manuscript. We further thank Kaizhen Li for the help with histology and immunolabelling, the electronic workshop and members of the Ciocchi Laboratory for their valuable inputs on the project. We thank Jakob Jordan and Nicolas Deperrois for fruitful discussions.

## Results

### An adjustable linear track maze to induce a scaling in anxiety behavior

The development of the aLTM allowed us to expose mice to 6 different anxiety configurations: at a height of 20 cm with the openable part closed (‘no anxiety’: NA) or open (‘very low anxiety’: VLA), with the openable part open at a height of 70 cm (‘low anxiety’: LA), 120 cm (‘moderate anxiety’: MA and ‘high anxiety’: HA) and 170 cm (‘very high anxiety’: VHA). Additionally, at the height of 120 cm, the floor width of the aLTM’s open part could be adjusted from 10 cm to 7 cm, resulting in two different configurations (i.e., MA and HA) while it remained with a 7 cm floor width at 170 cm height (Figure 1A, Materials and Methods). To ensure a comparable motivation and prevent habituation behavior to the aLTM’s open part across anxiety configurations, mice were food restricted and received small rewards (i.e., sugar pellets) in the aLTM’s openable/open part. At the behavioral level, we observed that the trials taken and the time spent by mice (N = 16) in the openable part (including successive higher or lower anxiety configurations) linearly correlated with the 6 anxiety configurations (simple linear regression: number of trials: *F* = 66.3, *R^2^* = 0.164, *p* < 0.001; percentage of time *F* = 38.3, *R^2^* = 0.102, *p* < 0.001; Spearman correlation: number of trials: *r* = -0.52, *p* < 0.001; percentage of time *r* = -0.36, *p* < 0.001) and consequently resulted in 6 behaviorally scaling anxiety levels: NA – VLA – LA – MA – HA – VHA (Figure 1B). A possible habituation/sensitization effect of the different anxiety configurations was further excluded by performing scrambled presentations of the different anxiety configurations, resulting in a linear scaling of behavioral anxiety when resorting the anxiety levels in an ascending order (Suppl. Figure 2). In classical anxiety tests such as the EPM, fewer entries and reduced time spent in the open arms compared to the closed arms have been shown to be related to higher anxiety (25, 32). Also in our task, we found a progressive reduction in trials taken and time spent within the aLTM’s open part, correlating with the increase in anxiety levels (Figure 1B, left and right respectively), and confirming that the anxiety configurations were inducing increasing anxiety in the open part of the maze.

### vH inhibition impairs the scaling of anxiety behavior

Inhibition of vH activity has been shown to reduce anxiety behavior (33-35). To explore the causal impact of vH activity in the aLTM, we applied an ArchT-mediated optogenetic intervention in vH CamKIIα^+^ neurons, thereby predominantly targeting excitatory pyramidal neurons, during the transition of mice (N = 6 mice per group) from the closed to the open part of the aLTM (Figure 2, Materials and Methods). With this approach, we aimed to influence vH activity primarily during the initial threat evaluation, and not during the complete exploration of the aLTM’s open part, to ensure that the perceived anxiety level the mice are facing is not masked by vH optogenetic manipulation. We found that the intervention in the transition zone only led to a significant increase in trials and increased time in the aLTM’s open part in the ArchT group of mice across anxiety-inducing configurations compared to the control mice (Figure 2D,E), indicating a reduction in anxiety. Moreover, inhibition of the vH during the transition to the aLTM’s open part disrupted the scaling of anxiety-related behavior expressed by unchanged trial numbers and time spent in the open part across anxiety levels in ArchT but not control mice (Figure 2F,G linear regressions of values of all sessions across all levels: *F* = 0.30, *R^2^*= 0.005, *p* = 0.586; *F* = 0.54, *R^2^* = 0.008, *p* = 0.465 for ArchT mice trials and time spent; *F* = 20.90, *R^2^* = 0.246, *p* < 0.001; *F* = 10.30, *R^2^* = 0.139, *p* = 0.002 for control mice trials and time spent. Spearman correlation: *r* = -0.13, *p* = 0.296; *r* = -0.13, *p* = 0.298 for ArchT mice trials and time spent and *r* = -0.49, *p* < 0.001; *r* = -0.35, *p* = 0.004). This suggests that interference with vH activity can impair the perception of anxiety levels.

### vH activity scales with increasing anxiety levels

Tetrode implantations within the vH enabled the recording of 207 single-units as mice navigated through the different anxiety configurations of the aLTM (Suppl. Figure 1). Previous studies have shown that the vH exhibits increased activity in open anxiogenic compartments of mazes (26, 27, 36, 37). Here, we show that individual neurons and vH population activity not only increased their activity in the open anxiogenic compartments, overrepresenting the anxiogenic part of the maze, but that the activity progressively increased from low to high anxiety configurations in the openable/open part of the maze (Figure 3). We analyzed vH activity across NA to HA configurations (VHA was not included due to insufficient open part exploration needed for reliable activity analysis) whereby each level was represented by activity originating from increasing and decreasing anxiety level presentations (see Materials and Methods). Simple linear regression analysis showed that the activity progressively increased from the NA to HA configuration in the openable/open part but not in the closed part in two single neuron examples [F = 29.39, *R^2^* = 0.406, *p* < 0.001 (Figure 3A left) and F = 64.80, *R^2^*= 0.601, *p* < 0.001 (Figure 3A right) for the openable/open part; *F* = 0.64, *R^2^* = 0.027, *p* = 0.432 (Figure 3A left) and F = 0.53, *R^2^* = 0.023, *p* = 0.474 (Figure 3A right) for the closed part] and the same observation could be made in the entire recorded vH population correlation values of: *F* = 43.00, *R^2^* = 0.040, *p* < 0.001 and *F* = 0.49, *R^2^*= 0.001, *p* = 0.485 for openable and closed part, respectively (Figure 3B). Additionally, the scaling of vH activity was not predominantly dependent on locomotion (Suppl. Figure 3).

### Tuning and reorganization of vH neuronal activity underlies scaling during anxiety

We hypothesized that two phenomena at the single neuron level could contribute to the increased and scaled neuronal activity that we observed in the aLTM’s open/openable part: first, a scaled activity tuning in which neurons could increase their activity with increasing anxiety levels and second, a neuronal recruitment to the open part of the aLTM driven by different anxiety levels. To reveal if such a process might be contributing to the scaled representation of anxiogenic information in the vH population activity, we selected neurons that increased their activity linearly (significant Spearman correlation, see Materials and Methods) across each anxiety configuration, i.e. from NA to VLA, from VLA to LA, from LA to MA and from MA to HA, in the open part of the maze. We found 30 neurons (14.5% of the total population) whose firing rates increased consecutively across the anxiety levels in the aLTM’s open but not the closed part and thereby showed increased tuning with increasing anxiety (Figure 4A, simple linear regression values: *F* = 52.60, *R^2^* = 0.150, *p* < 0.001 for openable, *F* = 0.51, *R^2^* = 0.002, *p* = 0.478 for closed).

To investigate if changes in neuronal recruitment took place at any specific anxiety configurations, we quantified the neuronal activity based on the earliest significant increase in firing in the aLTM’s open part compared to the closed part. Our results suggest that at each anxiety level, new neurons were recruited and maintained their activation to subsequent higher anxiety levels (Figure 4B-E). Although major reorganization occurred at the first exposure to the aLTM’s open part (in VLA), sequential recruitment occurred at any anxiety levels. This suggests that both enhanced tuning of neuronal responses and increased neuronal recruitment contribute to the scaling of neuronal activity in response to rising anxiety levels.

### Novelty does not change vH population activity

To discern changes induced by novelty from the changes related to anxiety configurations, we replaced the openable compartment with a novel compartment where the sidewalls had a clearly different visual appearance and a different texture (Figure 5A, right). During the exploration in the novel configuration, mice demonstrated a preference for the novel section of the maze (Figure 5B, left), but showed no difference in trials taken compared to the control configuration (corresponding to a NA configuration, Figure 5B, right). This shows that mice expressed a similar motivation to run trials and were able to detect and engage with the novel part of the maze (38).

Novelty can induce a spatial remapping of activity in the hippocampus (39, 40), but we observed that the novelty in the openable compartment, at least at the overall population level, did not lead to an increased vH activity compared to the control configuration (Figure 5C), suggesting that changes in vH activity during aLTM navigation were mainly due to anxiety processing.

### Decoding of anxiety levels by a convolutional classifier confirms the scaling of anxiety-related information

Finally, to further verify that vH population activity encodes anxiety-related information, and that this encoding scales with the actual perceived anxiety level, we measured the performance of a non-linear classifier trained to decode anxiety levels from recorded firing rates. The test accuracy on held-out data is above chance level for all configurations (Figure 6A), which is a telltale sign that vH population activity encodes information about anxiety levels. Next, to verify that this encoding scales continuously with each anxiety level, we leveraged the performance of the classifier on unseen data. For a classifier trained solely on normalized firing rate data from the two extreme conditions (i.e. NA and HA), the proportion of unseen data that are classified as NA (respectively as HA) can be interpreted as a measure of their similarity with the NA (respectively HA) configuration. For the vH to function as an anxiometer, we would thus expect the proportion of data being classified as NA to be the highest for actual NA data, and to continuously decrease as the actual anxiety level goes to VLA, LA, MA, and ultimately HA. This is indeed what we observed (Figure 6B), where the classifier was trained to solely discriminate NA and HA samples with its validation accuracy was assessed on held-out data. The leftmost value indicated the proportion of NA samples which were correctly classified as NA configurations. Similarly, the rightmost value indicates the proportion of HA samples which were classified as NA configuration. Expectedly, the proportion of samples classified as NA is decreasing when the ground truth anxiety level increases. Then, we assessed how unseen VLA, LA and MA samples would be sorted by the classifier only trained on NA and HA samples (second, third and fourth values). Overall, VLA samples have a higher probability of being classified as NA as compared to LA samples, which highlights a closer proximity between the VLA and NA configurations than between LA and NA. MA samples have a higher probability of being classified as HA as compared to LA samples, which highlights a closer proximity between the MA and HA configurations than between LA and HA. The proportion of unseen data that are classified as NA decreases with increasing anxiety level (Figure 6B, x-axis), which defines a continuously scaling ‘anxiety axis’ in the vH population activity. Simple linear regression analyses confirmed the observed scaling phenomenon with *F* = 31.72, *R^2^* = 0.227, *p* < 0.001. Collectively, the decoding of anxiety levels confirms the scaling of anxiety-related information.

**Figure 6.**
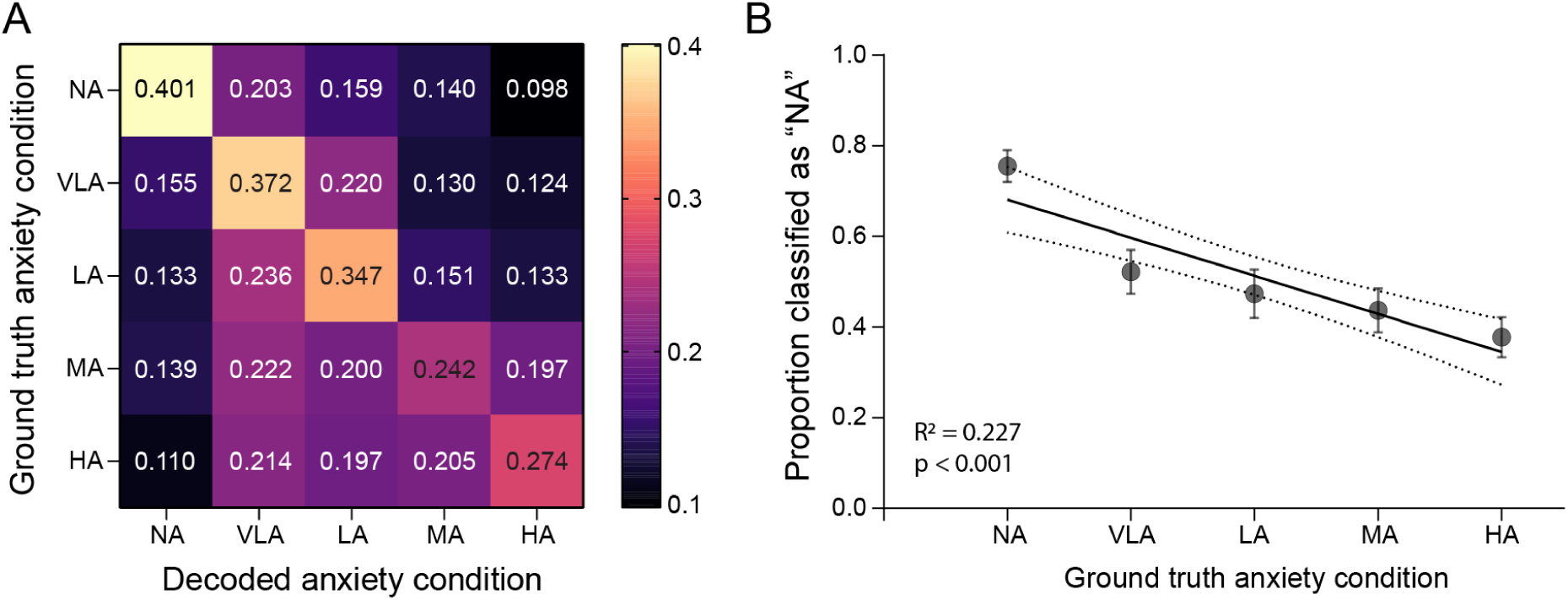
Decoding of anxiety levels by a convolutional classifier. (A) Confusion matrix indicating the validation accuracy. Chance level is 0.2 (since all 5 anxiety configurations have the same number of samples). **(B)** Convolutional classifier trained on NA and HA configurations for each experiment: held-out normalized firing rate data for these configurations are used as a validation set (HA and NA values) and shows that NA and HA configurations are correctly classified. The test set consists of all data from the VLA, LA and MA configurations (VLA, LA, MA values). Results show that the decoder output monotonically decreases as the anxiety level increases. Data represented as linear regressions with 95% confidence intervals calculated based on all behavioral sessions (N = 6 mice, n = 22 sessions). Means ± SEM of all behavioral sessions for each anxiety configuration are also indicated.

## Discussion

Humans and animals must strike a balance between minimizing exposure to possibly harmful events and exploring the environment to find rewards. Anxiety has been suggested to bias decision-making by increasing the prediction of negative future outcomes (41, 42). However, how anxiety coding in the brain is involved in risk assessment to guide behavior is not yet well understood. In our study, we investigated how anxiety representations in the vH of mice are tuned to different levels of anxiety and might thereby provide a basis for decisions under negative emotional conflict. To do so, we developed a maze, the aLTM, on which mice could be exposed to 6 different anxiety configurations during navigation in the same spatial environment while we recorded the activity of vH neurons.

We found that we could successfully induce different anxiety levels in mice and that anxiety-related activity in the vH progressively scaled with the different anxiety levels. Human neuroimaging studies have shown that greater hippocampal BOLD signals in the anterior hippocampus are associated with a greater probability of threat and higher anxiety during emotional conflicts and safety learning (21, 43). In rats, vH activity in the closed arm has been shown to be related to the extent to which animals explore the open part of a linear maze (31) and vCA1 activity has been shown to further increase with anxiety-associated head-dipping behavior (26). With our results, we provide evidence that behavioral scaling of anxiety within the same spatial and behavioral framework is supported by fine tuning and scaling of vH activity, suggesting that the vH might act as an ‘anxiometer’ in the brain. In contrast, previous work has shown that the basal amygdala encodes behavioral states such as freezing, head dips, entries and exits to anxiogenic compartments, but does not represent the level of absolute perceived anxiety (44). Rather, our findings position the vH as a critical limbic region that functions as an ‘anxiometer’ by scaling its activity based on perceived anxiety levels. To test the causal relationship between the scaling vH activity and anxiety experienced by the animals, we applied ArchT-mediated optogenetic intervention during the approach of the open compartment of the maze. In a study in humans, the decision to approach a reward relied on the hippocampus and not on the amygdala (20). In line with this finding, when we inhibited the vH during the approach of the open anxiogenic environment, the mice expressed reduced anxiety and a disruption in the scaling anxiety behavior in the open part of the maze. Hence, we posit that vH activity plays a crucial role not only in the generation of anxiety representations (45, 46), but our findings further substantiate that activity associated with anxiety, when facing anxiogenic situations, may instruct downstream areas to support executive function.

Tuning of neuronal activity has been extensively described in sensory areas, for example, in orientation-tuned neurons in the primary visual cortex that exhibit highest firing rates in response to specific orientation of a visual stimulus (47, 48). Responses of dopamine neurons in the ventral tegmental area of the midbrain have been also described to scale to error prediction depending on reward size (49). The integration of sensory stimuli such as odors, tones or speed of the animal have been shown in the hippocampus and entorhinal cortex to drive neural activity levels (50-53). How might individual neurons and population codes in higher cognitive areas such as the hippocampus achieve a tuning to internally generated emotional states such as anxiety? We described two processes that contributed to the scaling of anxiety-related activity in the vH, i.e. an enhanced tuning of neuronal responses and a neuronal recruitment to the open part of the aLTM driven by different anxiety levels. We hypothesize that these processes might be similar to the well described concepts of remapping in the hippocampus and rely on a spatial framework whereby neurons can either change their firing rates within the same location referred to as rate remapping or in a new environment randomly reconfiguring spatial codes during global remapping (54). We argue that these processes might not be restricted to the encoding of space but to any information that is predominantly processed by the underlying hippocampal subregion (55). The recruitment of new vH neurons that were activated at higher anxiety levels might reflect a process similar to global remapping whereby neurons that were not responding in previous anxiety encounters were engaged and recruited at a specific anxiety ‘threshold’, not to a new environment as during global remapping, but to a different anxiety level. vH circuits integrate a multiplicity of synaptic inputs: e.g., visual-spatial contexts and object information from the medial and lateral entorhinal cortex and anxiety-related information from the basolateral amygdala (34, 56, 57). We speculate that convergence of synaptic inputs and possibly inhibitory microcircuits might tune the scaling of rate remapping and gate the switching of newly recruited neurons during different anxiety encounters.

All changes of the anxiety configurations occurred within the same spatial environment, however the configurational changes, even minor, were of physical nature (e.g. distance to the ground or removal of part of the maze floor). To exclude that the mice might perceive these as novel aspects of the environment, we introduced a novel configuration of the maze. Based on our results, we conclude that even if a remapping had occurred, it did not result in such prominent dynamic increases of neuronal activity as observed across anxiety configurations and, therefore, novelty is unlikely to have driven the observed changes. The spatially independent tuning of activity supports the concept that the hippocampus integrates and represents emotional information based on perceived anxiety levels. To further support the notion of the vH acting as an ‘anxiometer’ at the populational level, we implemented a non-linear method for anxiety-scaling classification. After training a convolutional classifier on the two extreme anxiety configurations (no anxiety ‘NA’ and high anxiety ‘HA’), the probability of test data to be classified as NA can be interpreted as a measure of their similarity with the NA condition or HA condition respectively, and thus defines a linear “anxiety axis” going from NA to HA.

Anxiety disorders in humans are characterized by an elevated perception of potential threats leading to excessive uncontrollable worries that patients overcome with difficulties (58, 59). In patients with obsessive-compulsive disorders (a form of generalized anxiety) and schizophrenia (with frequent emotional disturbances), hyperactivity of the vH has been reported (60, 61). Also, patients with temporal lobe epilepsy and hyperexcitability of the vH can experience anxiety disorders (61, 62). We hypothesize that in some of these patients, the mechanisms to control and adjust the scaling of vH activity may be dysfunctional, possibly either shifted to more elevated levels or instead of gradually scaling rather jump to higher levels, maladaptively influencing their decisions and behavior. Collectively, our findings provide insight into how the hippocampus could allow for a dynamic and flexible adjustment of anxiety responses according to accumulated experience and prevailing environmental conditions. Adaptive anxiety scaling, when disrupted, may contribute to the manifestation of anxiety disorders.

---

**Suppl. Figure 1.**
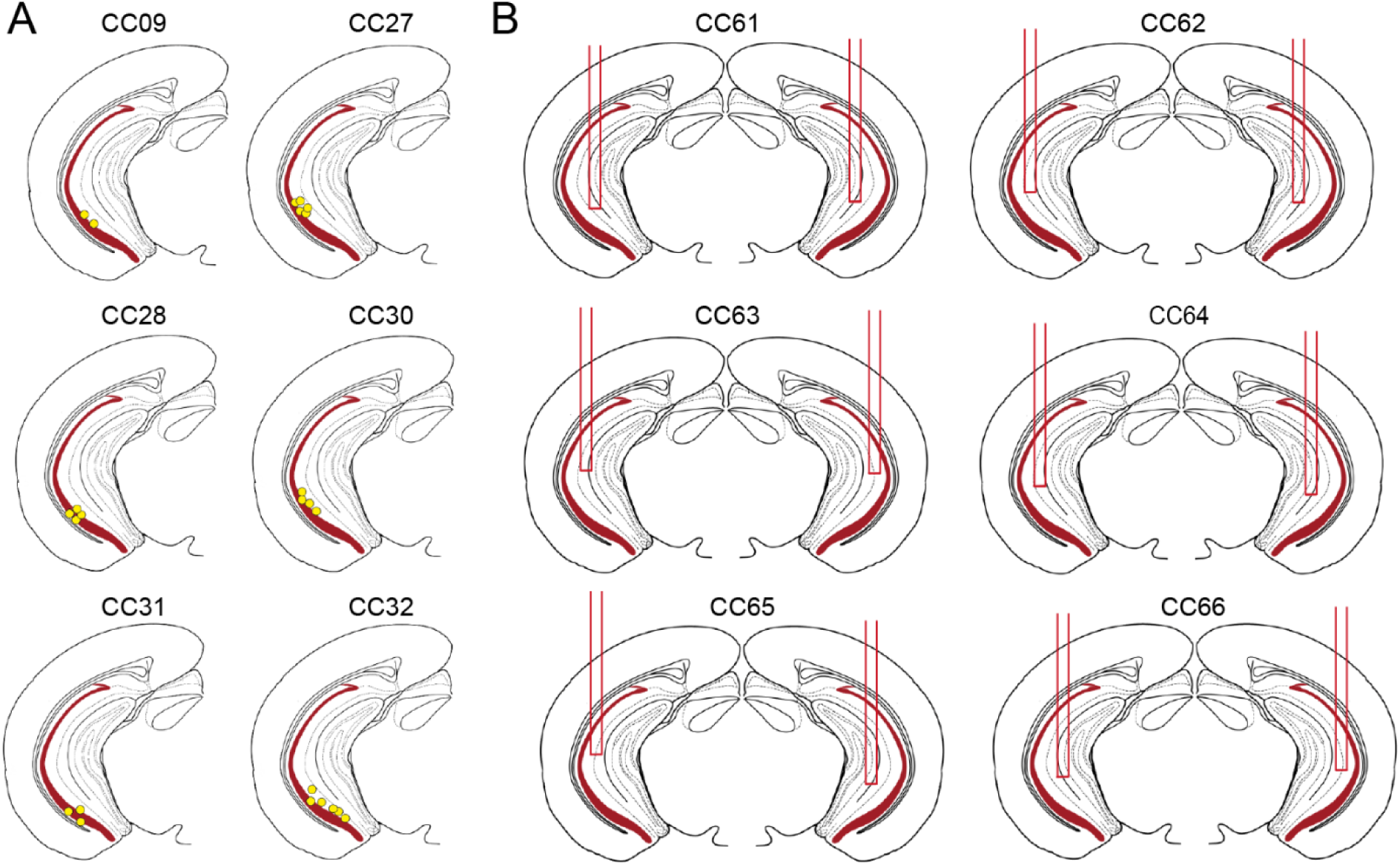
(A) Schematic representation of tetrode recordings sites indicated by electrolytic lesions (yellow circles). **(B)** Schematic representation of 6 optic fiber implants (red bars). Identifying animal names are on top of every schematic representation.

**Suppl. Figure 2.**
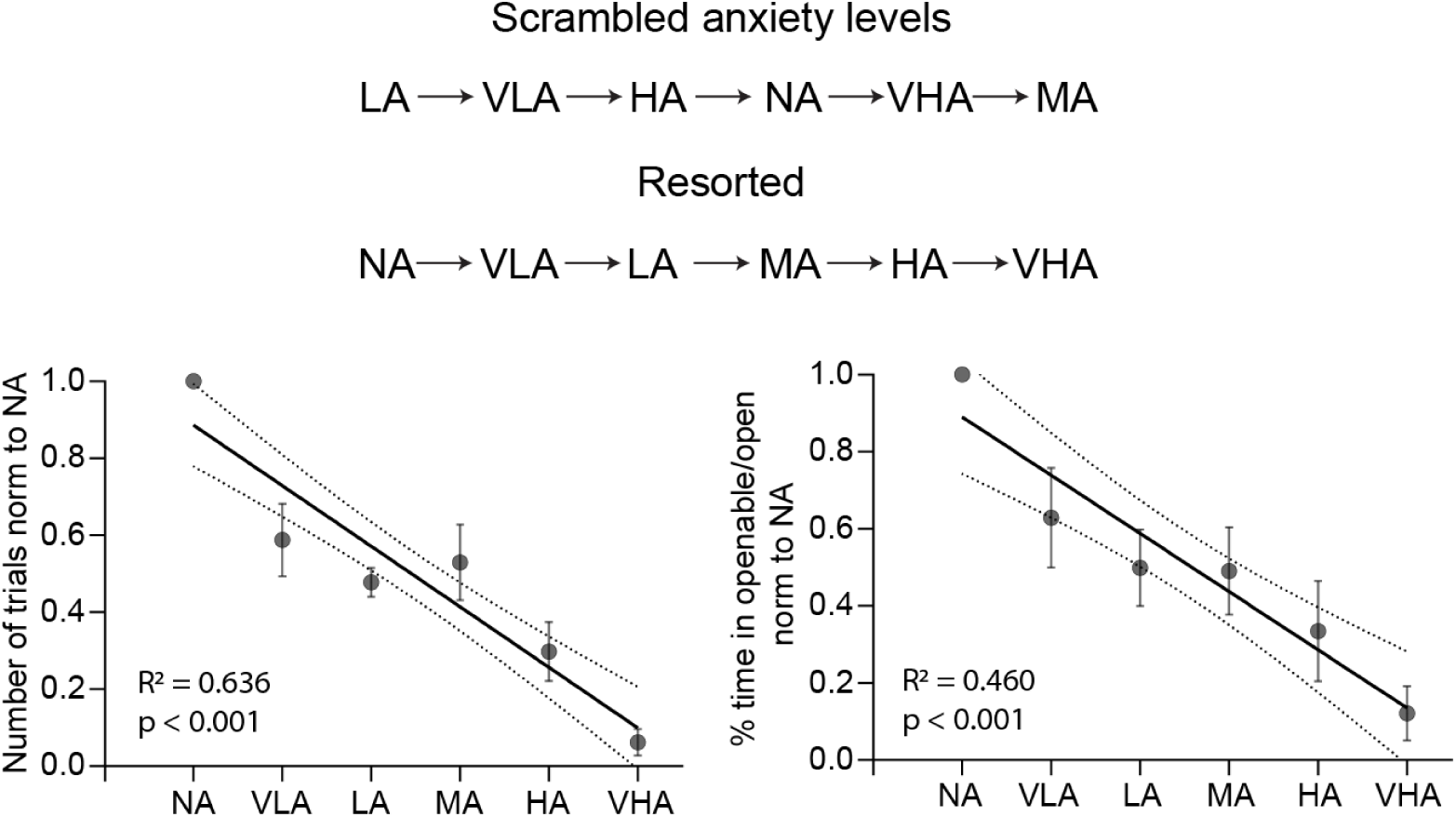
Scrambled control experiments, in which the sequence of presentation of the anxiety levels did not follow a linear order, provide evidence that mice can independently discriminate different anxiety levels on the aLTM. Number of trials (left) and percentage of time in open normalized to NA (right), simple linear regression model: *F* = 80.24, *R^2^* = 0.636, *p* < 0.001; *F* = 39.21, *R^2^* = 0.460, *p* < 0.001. Spearman correlation: number of trials: *r* = -0.80, *p* < 0.001; percentage of time *r* = -0.68, *p* < 0.001. For clarity reasons, the results are presented in resorted order (from NA to VHA). Data represented as linear regressions with 95% confidence intervals calculated based on all behavioral sessions (N = 8 mice, n = 48 sessions). Means ± SEM of all behavioral sessions for each anxiety configuration are also indicated.

**Suppl. Figure 3.**
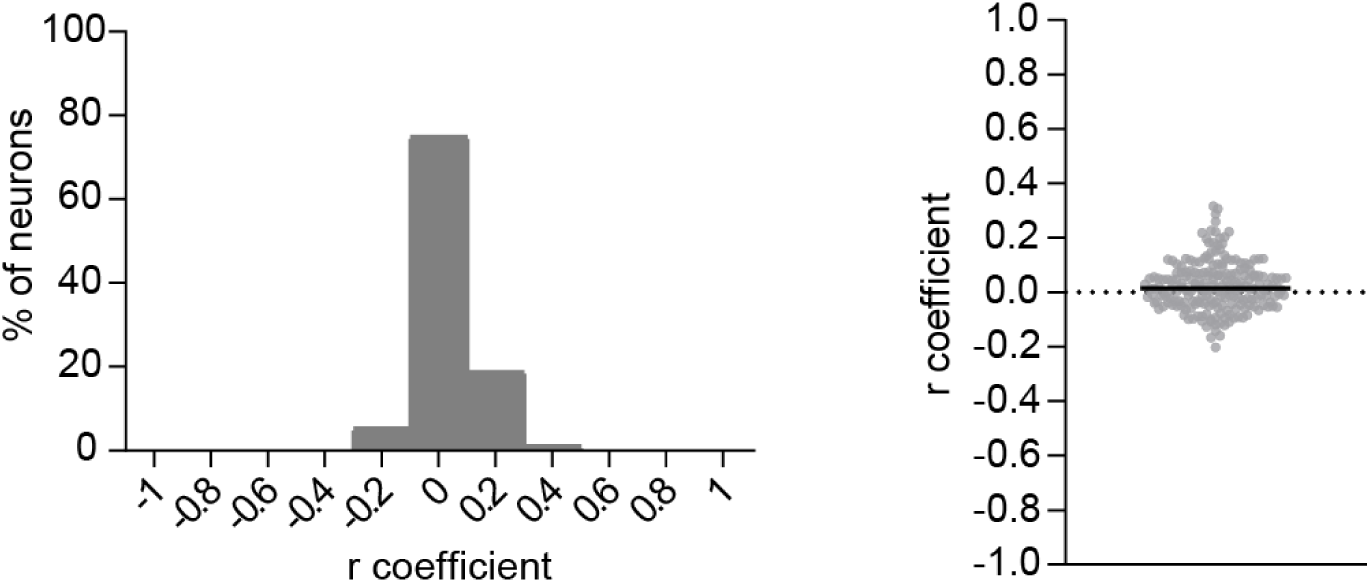
Correlational analysis of firing rates vs. speed during the exploration of the NA configuration. The majority of vH neurons present a *r* coefficient value close to 0 (left) indicating a small correlation between the firing rate and the speed of the mice. Distribution of individual *r* coefficient values plot (right) with a median of 0.008. N = 6 mice, n = 207 neurons.

